# Chemical signatures of honey bee group membership develop via a socially-modulated innate process

**DOI:** 10.1101/412353

**Authors:** Cassondra L. Vernier, Joshua J. Krupp, Katelyn Marcus, Abraham Hefetz, Joel D. Levine, Yehuda Ben-Shahar

## Abstract

Large social insect colonies exhibit a remarkable ability for recognizing group members via colony-specific cuticular hydrocarbon (CHC) pheromonal signatures. Previous work suggested that in some ant species colony-specific signatures are generated through a “gestalt” mechanism via the passive transfer and homogenization of CHCs across all individual members of the colony. In contrast, we demonstrate that nestmate recognition cues of worker honey bees (*Apis mellifera*) mature in foragers via a sequence of stereotypic age-dependent quantitative and qualitative chemical transitions, which are driven by intrinsic biosynthetic pathways. Therefore, in contrast to predictions of the “Gestalt” model, nestmate recognition cues in honey bee colonies do not represent a passive “average” signature that is carried and recognized by all colony members. Instead, specific colony members develop the relevant cues via an innately-determined developmental program that can be modulated by colony-specific social environmental factors.

## Introduction

The ability to recognize “self” plays an important role in regulating diverse processes across biological organizational levels (Tsutsui, 2004). Analogous to the acquired immunity system, which depends on self-recognition at the cellular and molecular levels (Boehm, 2006), adaptive organismal social interactions often depend on the ability of individuals to recognize group- and/ or genetic-relatedness of conspecifics to increase cooperation or to suppress inbreeding (Hamilton, 1964a, 1964b; Pusey and Wolf, 1996; Trivers, 1971; West et al., 2007; Wilkinson, 1988). One remarkable example of organismal recognition of “self” comes from colonies of social insects, which depend on a robust nestmate recognition system to prevent the loss of expensive resources to non-nestmates, and to maintain overall colony integrity (Hefetz, 2007; van Zweden and D’Ettorre, 2010).

As in other self-recognition systems, theoretical models suggest that nestmate recognition in social insect colonies depends on the ability of individual colony members to reliably match colony-specific phenotypic cues, or “labels”, carried by other colony members, to stored neural “templates” (Buckle and Greenberg, 1981; Errard, 1994; Gamboa et al., 1986; Getz, 1982; Hölldobler and Michener, 1980; Lacy and Sherman, 1983; Reeve, 1989; Tsutsui, 2004; van Zweden and D’Ettorre, 2010). In some social insect species, the cues used in recognizing individual members of the colony have been reported to be visual (Baracchi et al., 2015), but in most cases are thought to be chemical (van Zweden and D’Ettorre, 2010). These complex pheromonal cues are primarily composed of blends of cuticular hydrocarbons (CHCs) and fatty acids, which vary across colonies of the same species (van Zweden and D’Ettorre, 2010). Nevertheless, how these large groups of hundreds to thousands of individuals coordinate the production and recognition of a robust colony-specific chemical cue remains unknown for most species.

Because members of social insect colonies are often genetically related, it was initially assumed that the production of similar colony-specific pheromones by individual colony members is intrinsically driven by shared allelic variants (Crozier and Dix, 1979; Getz, 1982, 1981). However, empirical studies revealed that, surprisingly, genetic relatedness is not likely to play a major role in defining colony-specific cues relative to colony and/ or social environmental factors (Breed et al., 1988; Downs and Ratnieks, 1999; Heinze et al., 1996; Lahav et al., 2001; Liang and Silverman, 2000; Singer and Espelie, 1996; Stuart, 1988). Although these colony “environmental” factors remain unknown for most social insects species, it has been suggested that contributions from nest building materials (Breed et al., 1988; Couvillon et al., 2007; D’Ettorre et al., 2006; Espelie et al., 1990; Singer and Espelie, 1996), the queen (Carlin and Hölldobler, 1988, 1987, 1986, 1983), and diet (Buczkowski et al., 2005; Buczkowski and Silverman, 2006; Liang and Silverman, 2000; Richard et al., 2004, 2007) could, at least in part, provide specific chemical components to the chemical signature shared by all colony members. Consequently, empirical and theoretical studies have led to the development of a more widely accepted model, which states that individual colony members acquire their shared colony-specific chemical signature via a passive cue homogenization process, often referred to as the “Gestalt” model (Crozier and Dix, 1979). This model argues that over time, passive physical interactions between individuals and/ or between individuals and nest materials leads to the accumulation of an average chemical signature that is present in all colony members (Crozier and Dix, 1979). To date, empirical data in support of the “Gestalt” model have been reported for a few ant species, which are known to transfer mixed blends of CHCs between individuals through trophallaxis and grooming via the action of the postpharyngeal gland (PPG) (Boulay et al., 2000; Lenoir et al., 2001; Meskali et al., 1995; Soroker et al., 1994; Victoria Soroker et al., 1995; Van Zweden et al., 2010). More recently it has been suggested that the cephalic salivary gland might serve a similar function as the PPG in the honey bee (Martin et al., 2018). However, some ant species do not display trophallaxis or allogrooming behaviors, and data suggest that not all ant species exhibit a PPG-derived common chemical profile amongst colony members. Instead, in some ant species, individual chemical profiles differ as a function of genetic relatedness and age (Teseo et al., 2014). Therefore, whether the “Gestalt” model represents a general mechanism for defining chemical cue specificity in nestmate recognition systems across all social insect species remains unknown.

Because of its economic importance and long association with human agriculture, one of the best studied social insect species is the European honey bee, *Apis mellifera*. Numerous previous studies have demonstrated that honey bees exhibit a robust nestmate recognition system that is based on the chemical recognition of pheromones (van Zweden and D’Ettorre, 2010). Analyses of CHC profiles showed that newly emerged honey bee workers express significantly lower amounts of total CHCs and lower overall CHC chemical diversity in comparison to older foragers, which are expected to elicit the strongest nestmate recognition response from guards at the entrance to the hive (Breed et al., 2004; Kather et al., 2011). Based on these studies, it has been hypothesized that the CHC profile of newly eclosed workers is analogous to a “blank slate”, and that, similar to the “Gestalt” model, nestmate recognition cues are homogenized and redistributed to colony members through intermediate environmental factors (Breed et al., 2015, 2004). Although experimental data suggest that honey bee nestmate recognition cues likely rely upon environmental sources (Downs and Ratnieks, 1999), such as the honeycomb wax (Breed, 1998; Breed et al., 1988; Couvillon et al., 2007; D’Ettorre et al., 2006), whether honey bees acquire their CHC profile via a passive “Gestalt”-like mechanism has not been investigated.

Here we provide empirical evidence that honey bees have evolved a different mechanism for the expression of colony-specific nestmate recognition cues that does not seem to depend on the “Gestalt” model. Our data indicate that in contrast to predictions of the “Gestalt” model, the maturation of the pheromonal cues required for nestmate recognition by individual honey bee workers is primarily regulated by developmental processes associated with behavioral task and the colony environment. Specifically, we find that individual workers exhibit stereotypic quantitative and qualitative changes in their CHC profile as they transition from in-hive tasks to foraging outside, and that the foraging task is directly associated with the final maturation of the colony-specific nestmate recognition cue. Together, our findings suggest that, contrary to a stipulation of the “Gestalt” model (Crozier and Dix, 1979), not all members of honey bee colonies display a uniform chemical signature via the passive acquisition of CHCs. Instead, our data suggest that nestmate recognition cues in honey bees are more likely a product of an interaction between a genetically-determined developmental program and factors associated with the colony environment.

## Results

### CHC profiles of individual honey bee workers exhibit qualitative and quantitative age-dependent changes

Given that newly emerged honey bees have lower amounts of total CHCs, and exhibit less chemical diversity compared to older bees (Breed et al., 2004), we initially sought to determine the age at which the CHC profile of individual honey bee workers matures. To achieve this goal, we analyzed the CHC profiles of individual workers from a single age-cohort that was reintroduced into its natal colony and then collected at different ages. This revealed that the total amount of CHCs increases between one-day post-reintroduction and seven-days post-reintroduction and then remains stable (Figure 1A, Kruskal-Wallis, H = 9.21, df = 3, p = 0.026; Figure 1 – figure supplement 1A, ANOVA, F(3,28) = 6.40, p = 0.002). Independently of the age-related quantitative changes, we also found that the CHC profiles of workers exhibit age-related qualitative changes in the overall CHC chemical composition (Figure 1B, Permutation MANOVA, F(1,31) = 22.86, R^2^ = 0.43, p < 0.001; Figure 1 – figure supplement 1B, Permutation MANOVA, F(3,28) = 2.35, R^2^ = 0.22, p = 0.038), as well as in the relative amounts of individual CHCs (Figure 1C, D, Figure 1 – figure supplement 1C, D, Table 1, Table 1 – table supplement 1). These data suggest that age might be playing an important role in the regulation of both the quantitative and the qualitative dimensions of the cuticular chemical profile of individual honey bee workers. These data also indicate that, in contrast to an assumption of the “Gestalt” model, not all members of a honey bee colony share a common chemical signature.

**Table 1:**
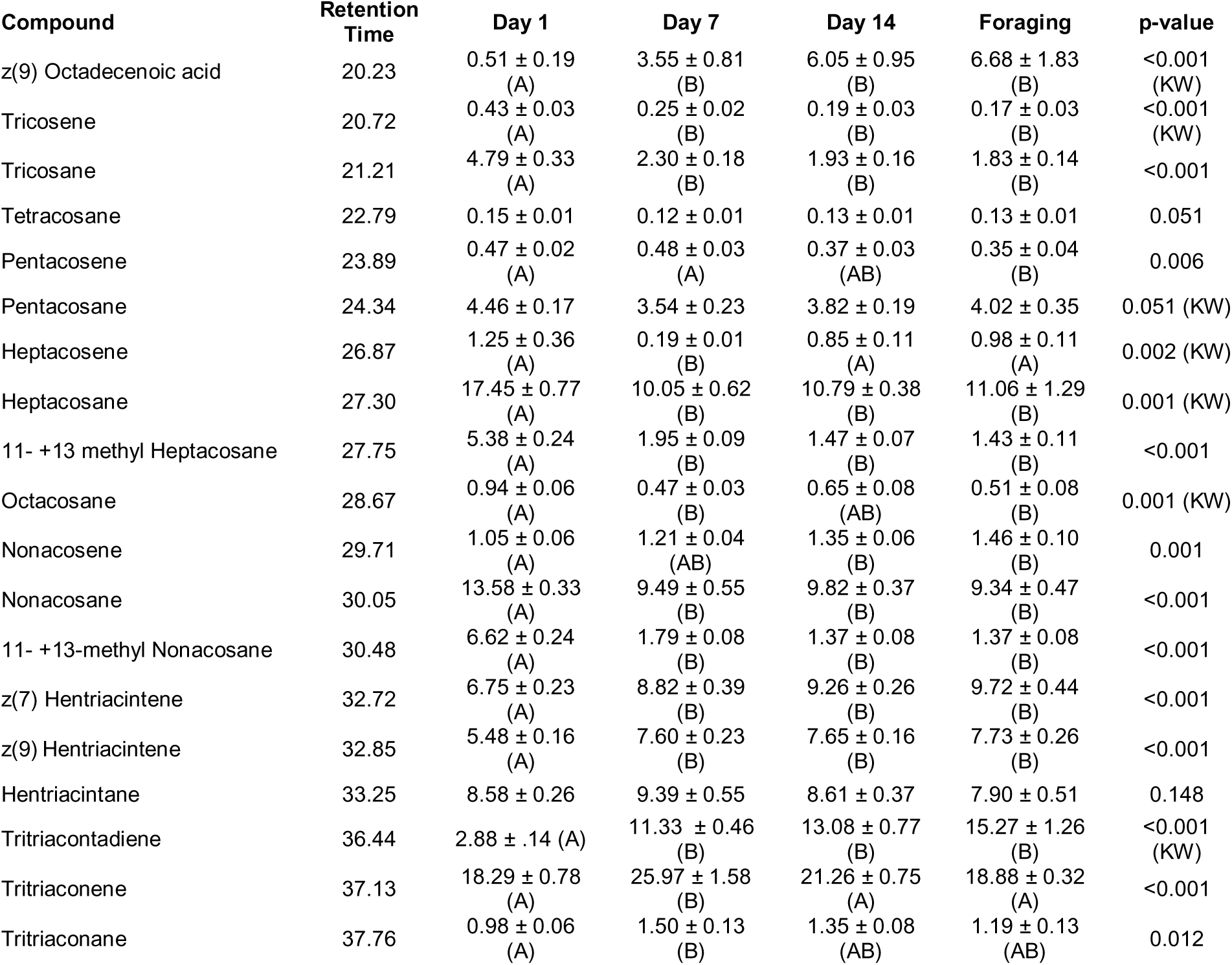
Individual compounds vary in proportion across different aged sister bees of a single colony. Numbers represent mean proportion of compound across bees of that age ± standard error. All p-values are from parametric ANOVA or nonparametric Kruskal Wallis ANOVA (denoted by “KW”). Letters denote statistically significant age groups across individual compounds via Tukey’s HSD (ANOVA post- hoc) or Dunn’s Test with FDR adjustment (KW post-hoc) (p < 0.05).

**Figure 1:**
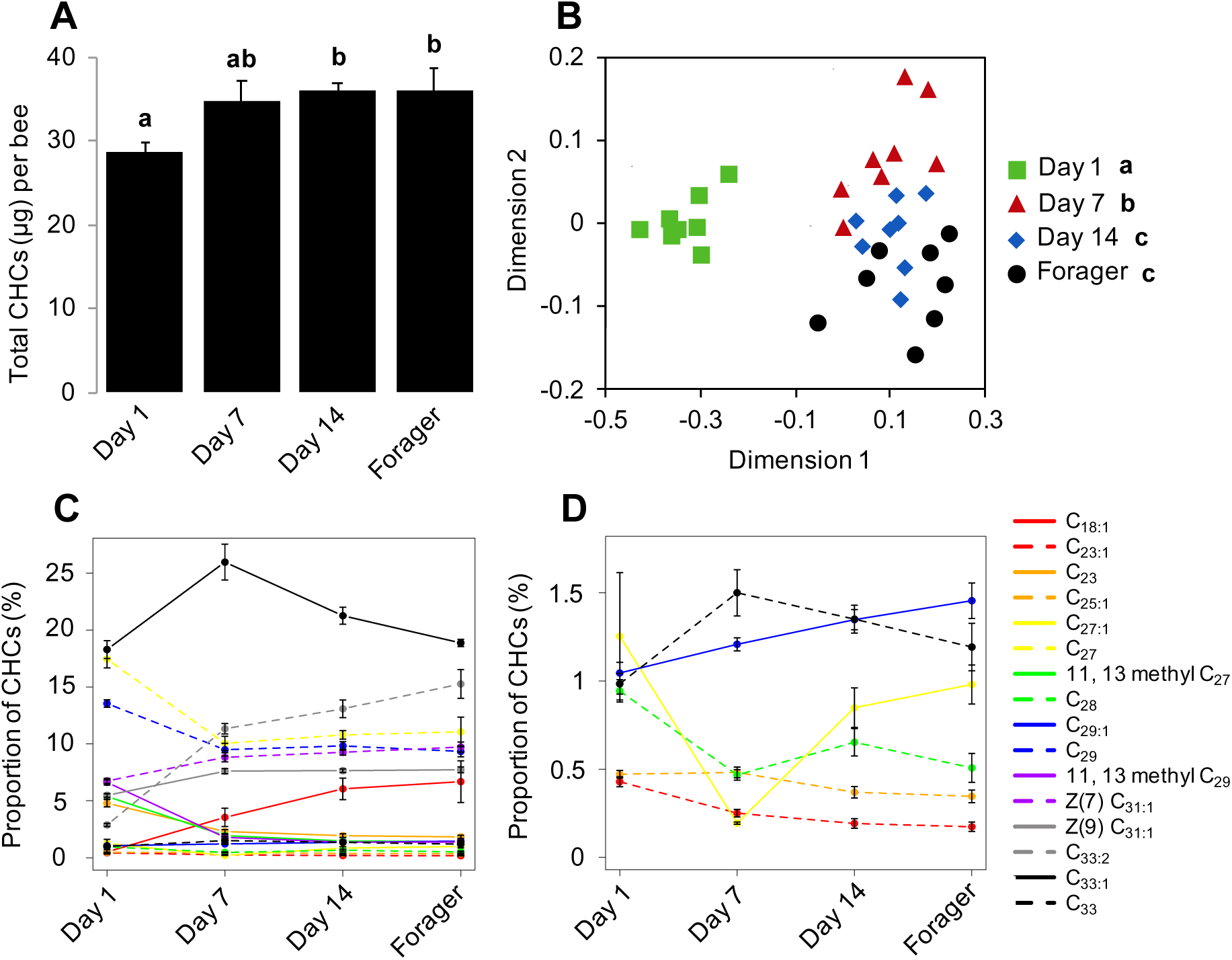
CHC profiles of bees exhibit quantitative and qualitative changes in association with age. (A) Total CHC amounts (μg) extracted from sister bees of different ages. (B) CHC profiles of sister bees of different ages. (C) Statistically significantly changing proportions of individual CHCs across sister bees of different ages. (D) A subset of C with low proportions. Statistics in A using ANOVA followed by Tukey’s HSD post-hoc. Statistics in B using Permutation MANOVA followed by FDR pairwise contrasts shown as a non-metric multidimensional scaling plot depicting Bray-Curtis dissimilarity between samples. Statistics for C & D are listed in Table 1. Lowercase letters above bars in A and legend in B denote *posthoc* significance (p < 0.05). Sample size per group, N = 8.

### The CHC signatures of individual workers are task-related

Honey bee workers exhibit age-related division of labor, which is characterized by a stereotypic sequence of in-hive behavioral tasks such as nursing and food handling, followed by the final transition to foraging outside the colony at about three weeks of age (Robinson, 1992; Smith et al., 2008; Søvik et al., 2015). Consequently, under natural colony settings, it is impossible to separate the possible independent impacts of ‘age’ and ‘task’ on the expression of forager-specific CHC profiles. Therefore, we next analyzed the CHC profiles of individual nurse and forager bees from single-cohort-colonies (SCC), a well-established experimental approach to uncouple behavioral maturation from chronological age (Ben-Shahar et al., 2004, 2002; Greenberg et al., 2012; Whitfield et al., 2003). Because these artificial colonies are initially comprised of a single age-cohort of day-old bees, a small proportion of these young workers will accelerate their behavioral maturation to become precocious foragers that are the same age as typical nurses (∼7 days old) (Ben-Shahar et al., 2002; Greenberg et al., 2012; Huang and Robinson, 1992). The comparison of the CHC profiles of typical young nurses and precocious foragers of identical age revealed a significant effect of task on the CHC profile of individual workers (Figure 2A, Permutation MANOVA, F(1,15) = 13.79, R^2^ = 0.50, p < 0.001). Similarly, we observed a significant effect of task on the CHC profiles of individual “over-aged” nurses and typical-aged foragers at three weeks of age (Figure 2B, Permutation MANOVA, F(1,15) = 45.41, R^2^ = 0.76, p < 0.001). In contrast, task and age had no effect on total CHC amount (Figure 2C, Two-way ANOVA, age: F(1,28) = 0.55, p = 0.46, task: F(1,28) = 0.37, p = 0.55, age*task: F(1,28) = 5.37, p = 0.03).

**Figure 2:**
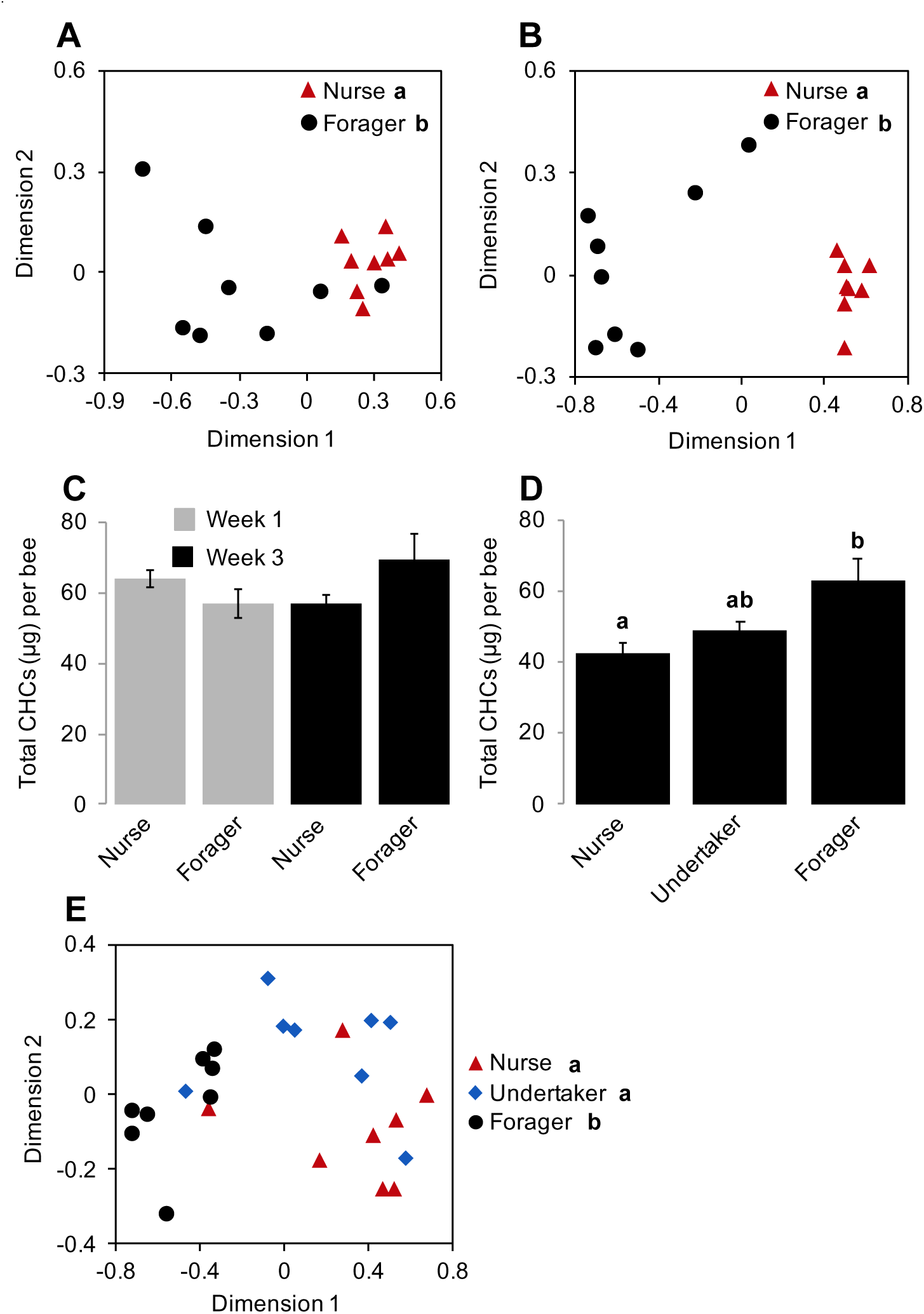
Effect of task on the CHC profile of bees is independent of age. Single cohort colony bees differ in CHC profile by behavioral task at one week of age (typical nurse age, (A)) and three weeks of age (typical forager age, (B)). (C) SCC bees do not differ in total CHC amount due to age and/or task. (D) Undertakers do not differ in total CHC amount from nurses and foragers. (E) Undertakers and nurses differ from foragers in CHC profile. Total CHC statistics (C, D) using ANOVA followed by Tukey’s HSD with FDR correction. CHC profile statistics (A, B, E) using Permutation MANOVA followed by FDR pairwise contrasts, shown as non-metric multidimensional scaling plots depicting Bray-Curtis dissimilarity between samples. Letters in graphs and legends denote *posthoc* statistical significance (p < 0.05). Sample size per group, N = 8.

Together, these data suggest that processes associated with the behavioral maturation of honey bee workers, not chronological age, are primarily responsible for the observed forager versus nurse CHC signatures of individual honey bee workers.

Previous studies suggested that exposure to the environment outside the colony is sufficient to induce changes in the CHC profiles of individual social insects (Wagner et al., 2001), which may account for our observed forager-specific CHC profiles in honey bees. Therefore, we next asked whether spending time outside the hive is sufficient to induce the observed foraging-specific CHC signature by comparing the CHC profiles between “undertakers”, nurses, and foragers from typical colonies. “Undertakers” are a small group of highly specialized older pre-foraging workers, which are responsible for removing dead bees by carrying them outside and away from the colony (Robinson, 1992; Smith et al., 2008; Søvik et al., 2015). Therefore, because undertakers and foragers perform their respective tasks outside the hive, while nurses and other younger, pre-foraging bees rarely do, we reasoned that if outdoor exposure defines the distinct forager-specific CHC signatures then the CHC profiles of undertakers should be more similar to foragers than to nurses. However, we found that although undertakers do not differ from foragers in total CHC amount (Figure 2D, ANOVA, F(2,21) = 6.228, p = 0.008), their CHC profiles are qualitatively markedly different from foragers, yet similar to those of nurses (Figure 2E, Permutation MANOVA, F(2,23)=12.60, R^2^ = 0.55, p < 0.001). These data suggest that the forager-specific CHC signatures are not solely driven by exposure to the outside environment, and therefore, are likely driven by the worker’s physiological transition to foraging activity.

### The development of individual CHC profiles is a regulated process modulated by the colony environment

Previous work indicates that guard bees will accept foraging- age nestmates and reject foraging-age non-nestmates, independent of genetic relatedness (Downs and Ratnieks, 1999). This suggests that factors associated with the hive environment play a dominant role in specifying the colony-specific chemical signatures used for nestmate recognition. Yet, our data also indicate that CHC development in individual workers is a developmentally-regulated process that is closely associated with the age-dependent division of labor among workers. To address this potential conundrum, we next asked whether the effects of task and colony environment on the development of CHC profiles of individual workers are independent by using a reciprocal cross-fostering strategy. To achieve our goal, we introduced cohorts of newly eclosed bees from two different typical colonies back into their own natal colony, as well as a reciprocal foster colony, and then recollected marked workers from both cohorts in each reciprocal colony at different ages. CHC analyses revealed that through Day 14, the CHC profiles of fostered bees were more similar to the profiles of their natal sisters than those of the host bees of the same age (Figure 3A, Permutation MANOVA, F(3,31)=3.15, R^2^ = 0.25, p < 0.001; Figure 3B, Permutation MANOVA, F(3,31)=2.02, R^2^= 0.18, p = 0.04). In contrast, once workers shift to foraging activity, we found that the CHC profiles of fostered bees are different from the profiles of both foraging natal sisters and unrelated host foragers of similar age (Figure 3C, Permutation MANOVA, F(3,31) = 4.06, R^2^ = 0.30, p = 0.002). Together, these data suggest that factors associated with the natal colony play an important role in defining the CHC profile of individuals during the early phases of behavioral development. However, by the time bees start foraging, the mature CHC profile of individual workers is defined by an interaction between factors associated with both the natal and foster colonies (Figure 3C).

**Figure 3:**
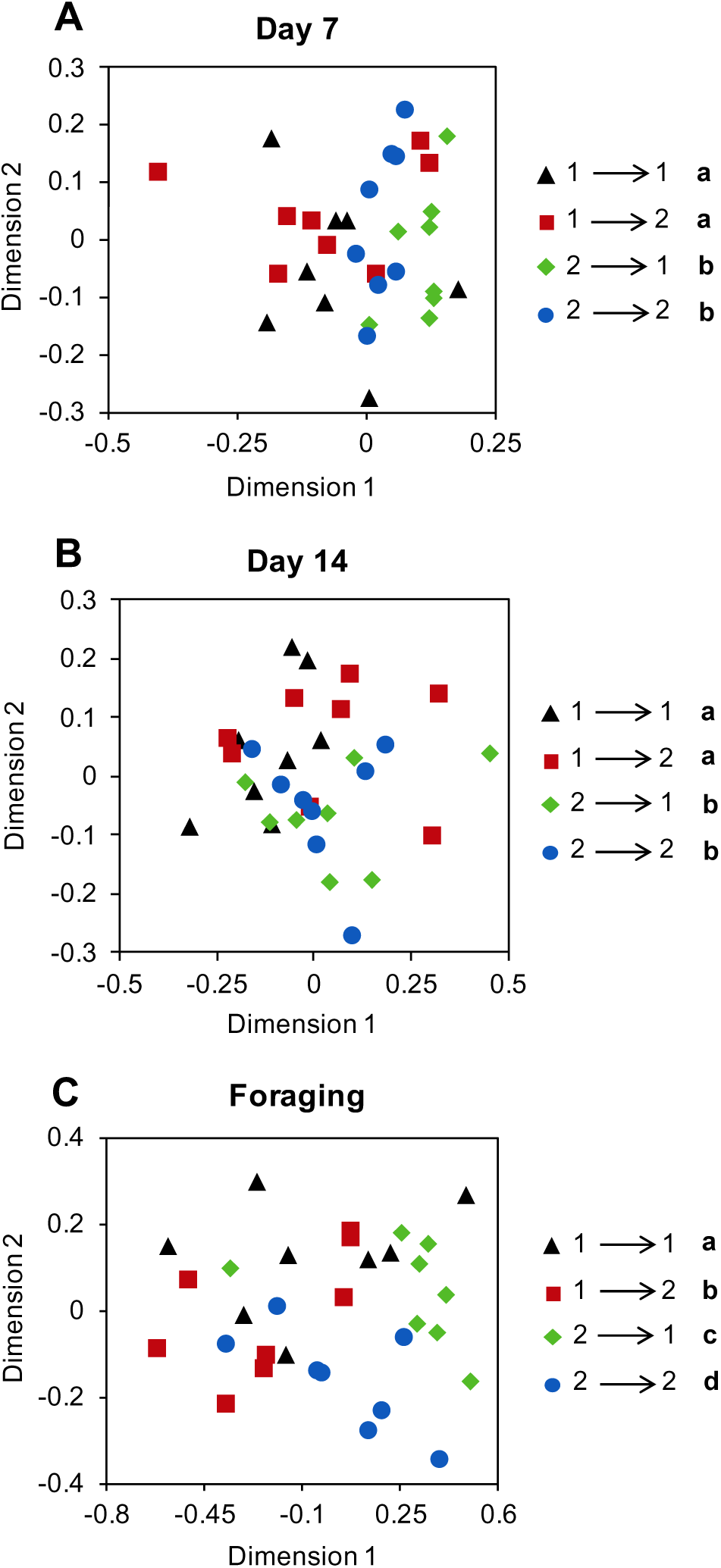
Cross-fostering indicates colony environment manifests in the CHC profile just prior to foraging behavior. Age-matched cross-fostered bees differ in CHC profile by natal colony at Day 7 (A) and Day 14 (B), and by both natal colony and foster colony when they are foragers (C). Number to left of arrow in legend represents the bee’s natal colony, and the number to the right represents the bee’s foster colony. All statistics using Permutation MANOVA followed by FDR pairwise contrasts, shown as non-metric multidimensional scaling plots depicting Bray-Curtis dissimilarity between samples. Letters in legends denote *posthoc* statistical significance (p < 0.05). Sample size per group, N = 8.

### The development of CHC profiles of individual workers is associated with the regulation of CHC biosynthesis genes

The “Gestalt” model predicts that nestmate recognition cues are passively acquired by individuals via physical contact with other colony members and/or environmental sources of hydrocarbons (Crozier and Dix, 1979; Lenoir et al., 2001; Meskali et al., 1995; Soroker et al., 1994; Victoria Soroker et al., 1995; Van Zweden et al., 2010). However, because our data indicate that the maturation of the CHC profile of individual honeybee bees is actually regulated in association with age-dependent division of labor, we next hypothesized that, in contrast to the passive “Gestalt” model, the CHC profiles of worker honey bees develop, at least in part, via active physiological and molecular task-related processes in the oenocytes, the abdominal subcuticular CHC producing cells of insects (Chung and Carroll, 2015; Falcón et al., 2014; Makki et al., 2014; Yew and Chung, 2015). We tested our hypothesis by examining whether age and/or task have an effect on the mRNA expression levels of genes that encode elongases and desaturases, the major enzymes in the CHC biosynthesis pathway (Chung and Carroll, 2015). To achieve this, we first used a bioinformatic approach to identify all putative members of both protein families in the honey bee genome (Figure 4). Next, we used real-time quantitative RT-PCR to compare mRNA levels of each candidate gene in abdominal cuticles from bees of different ages raised in their natal colony, as well as between foraging sister bees raised in either their natal colony or an unrelated foster-colony. Our analyses revealed that the expression of several elongases and desaturases is regulated by either age or colony environment (Figure 5 and Figure 5—figure supplement 1). Together, these data further indicate that, in contrast to the assumption of the “Gestalt” model and previous reports on how some ant species acquire their nestmate recognition cues, the CHC profiles of individual honey bee workers do not represent an average cue that is passively acquired via physical interactions between individuals or between individuals and nest materials. Instead, the CHC profile of individual honey bee workers is likely determined by an innate developmental process that could be modulated by factors such as age-, task-, and colony environment.

**Figure 4:**
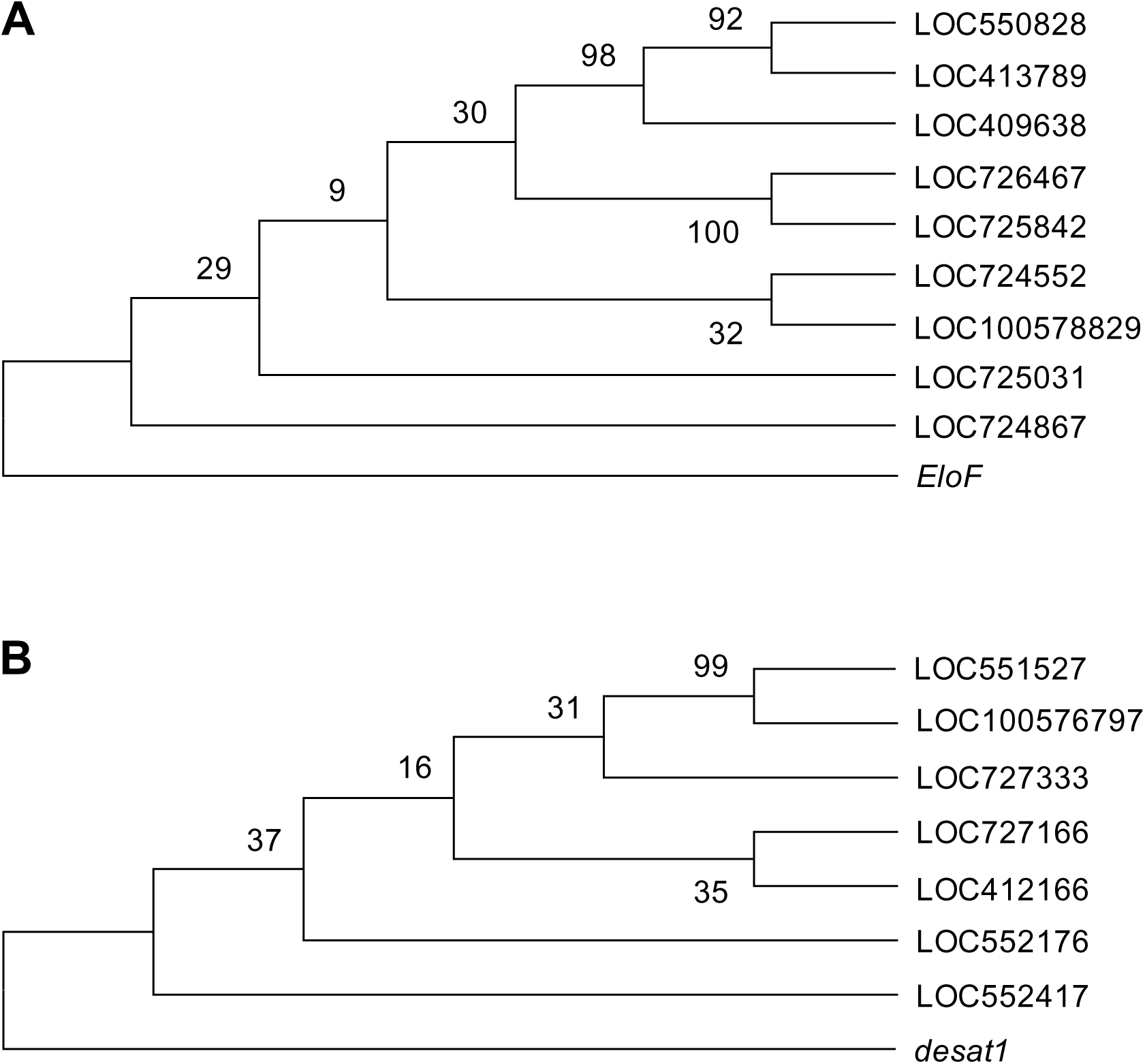
Elongase and Desaturase protein families in the honey bee genome. (A) Maximum likelihood bootstrap consensus tree of honey bee elongase genes with the *D. melanogaster* gene *EloF* as an outgroup. (B) Maximum likelihood bootstrap consensus tree of honey bee desaturase genes with the *D. melanogaster* gene *desat1* as an outgroup. Numbers next to branches represent the percentage of replicate trees, out of 500 replicates, in which the associated taxa clustered together.

**Figure 5:**
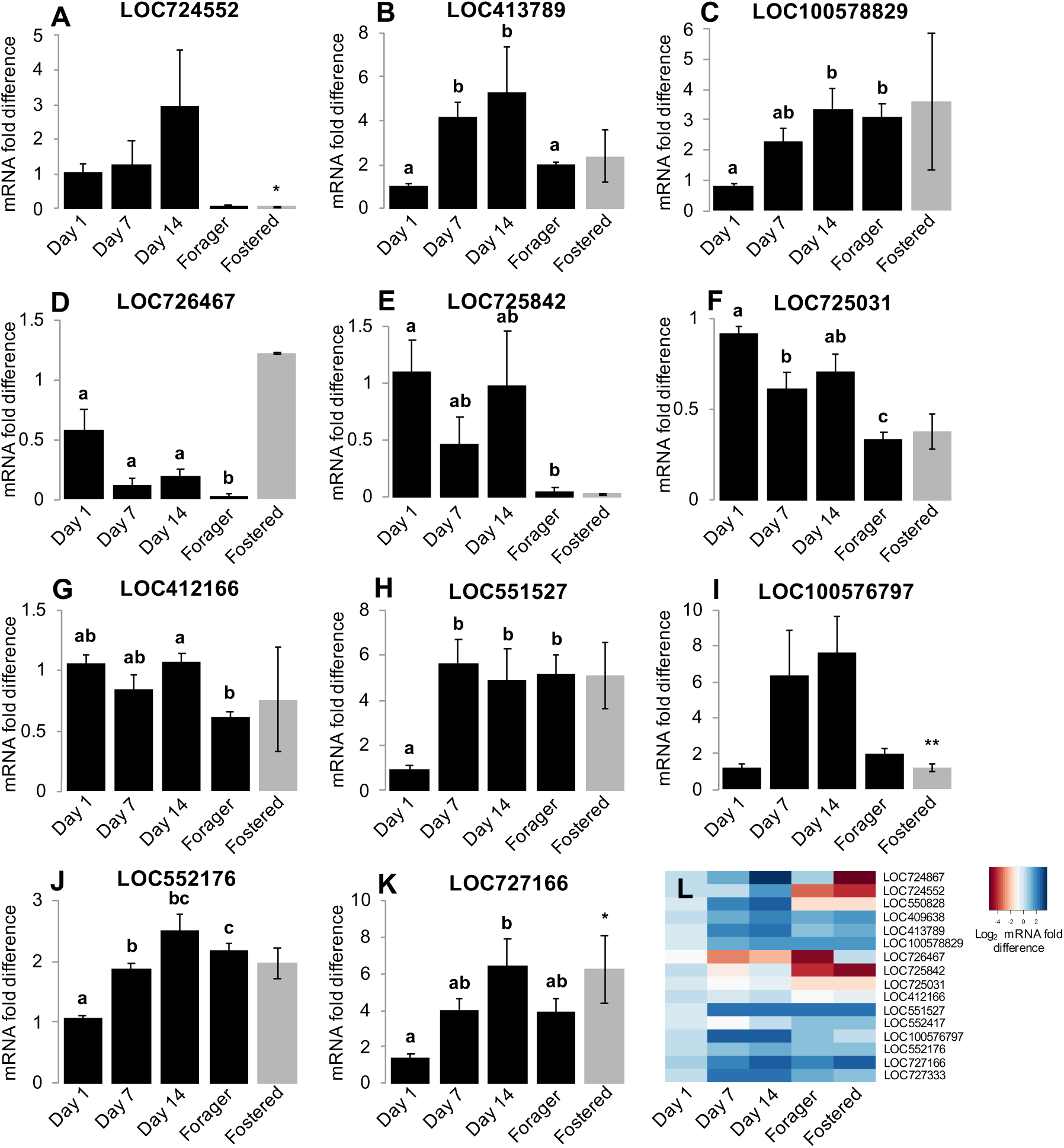
Age and social environment affect the expression level of CHC biosynthesis genes. (A-F) Elongase genes. (G-K) Desaturase genes. Only genes that different expression levels between at least two groups are shown (See Figure 5 – figure supplement 1 for results for all studied genes). Black bars represent bees raised in their own colony. Grey bars represent forager bees that were raised in an unrelated colony (“Fostered”). (L) Heat map of relative expression levels of all genes tested. Aging bee statistics using ANOVA followed by Tukey HSD post-hoc, or Kruskal-Wallis followed by Dunn’s Test with FDR adjustment post-hoc, with letters denoting *posthoc* statistical significance (p < 0.05). Between colony statistics using Mann-Whitney U test, with asterisks above grey bars denoting statistical significance from foraging bees raised in their own colony (*, p < 0.10; **, p < 0.05). Sample size per group, N = 4.

### Age and task play a role in defining nestmate recognition cues in honey bee colonies

Together, our data suggest that, in contrast to some of the assumptions of the “Gestalt” model, it is unlikely that all members of a single honey bee colony share the colony-specific nestmate recognition cues. Instead, our data support a different model, which stipulates that the specific chemical cues used for nestmate recognition develop in association with the well-described age-dependent division of labor in this species (Robinson, 1992; Smith et al., 2008; Søvik et al., 2015), reaching maturation just prior to the workers’ final transition to foraging behavior. To test this hypothesis, we investigated the behavioral responses of related and unrelated guard bees to focal bees of different ages (Day 1, Day 7, Day 14, and foragers on Day 21). At each test colony, the response of guards to random returning foragers from the natal and an unrelated colony were used as the benchmark for the baseline level of nestmate recognition behavior. Our data indicate that bees are accepted at their own colony, regardless of age (Figure 6A, Pearson’s Chi-Squared, Day1: χ^2^ = 49.05, df = 2, p < 0.001, Day 7: χ^2^ = 19.07, df = 2, p < 0.001, Day 14: χ^2^ = 44.89, df = 2, p < 0.001, Day 21: χ^2^ = 28.32, df = 2, p < 0.001). In contrast, at an unrelated colony, bees were accepted on Days 7 and 14, but rejected as foragers (Day 21) (Figure 6B, Day1: χ^2^ = 11.61, df = 2, p = 0.003, Day 7: χ^2^ = 15.51, df = 2, p < 0.001, Day 14: χ^2^ = 11.91, df = 2, p = 0.002, Day 21: χ^2^ = 7.35, df = 2, p = 0.04). These data further support the idea that colony-specific cues are likely not carried by honey bees until shortly before they begin foraging.

**Figure 6:**
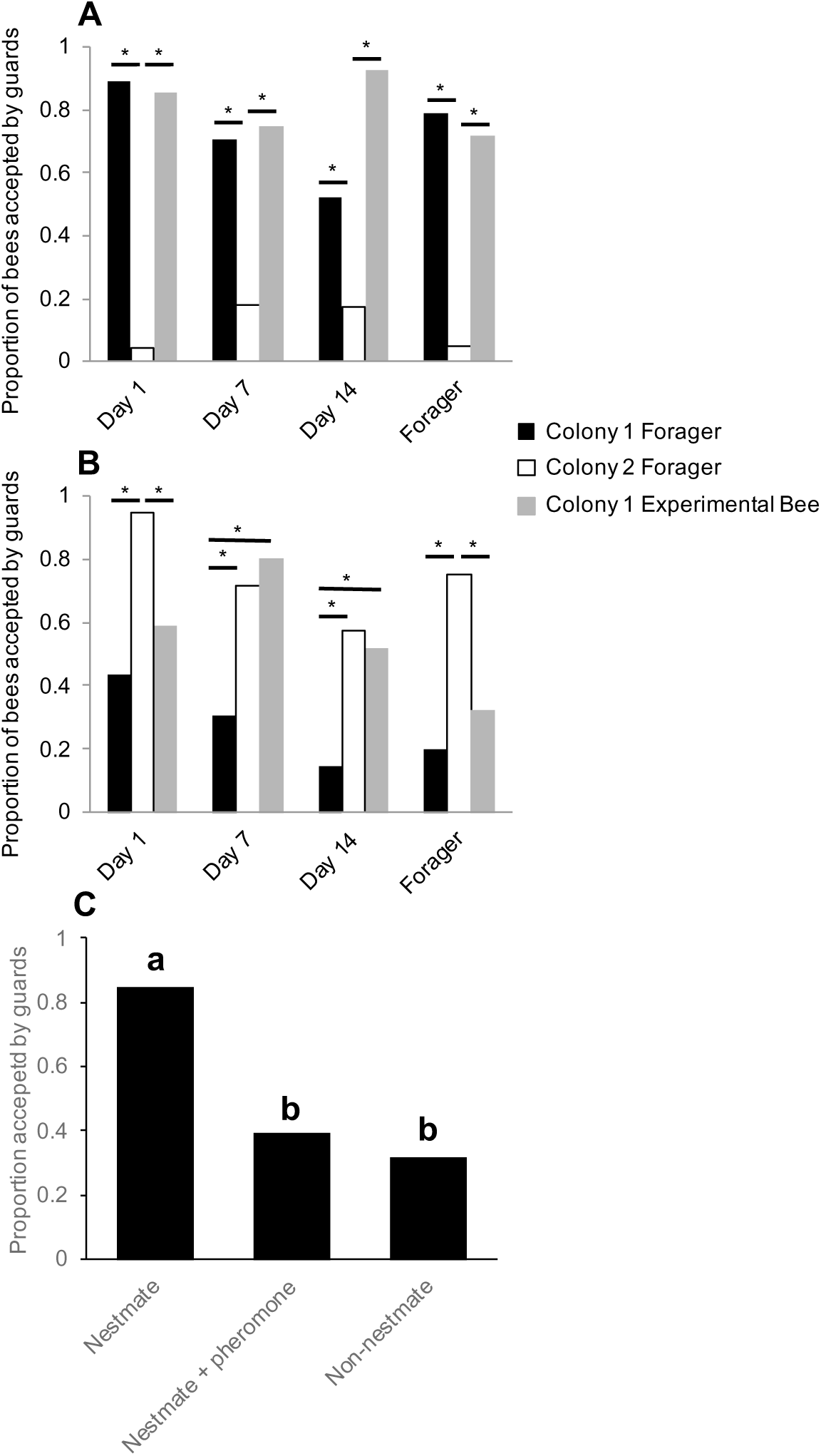
Bees develop their nestmate recognition cues shortly before becoming foragers. (A) Bees are accepted at a similar rate as Colony 1 foragers at the entrance to their natal colony (Colony 1) at all ages. (B) Bees are rejected at a similar rate as Colony 1 foragers at an unrelated colony (Colony 2) on Day 1 and Day 21. However, bees are accepted at a similar rate as Colony 2 foragers at an unrelated colony (Colony 2) on Day 7 and Day 14. (C) Nestmates are rejected by guards under acute exposure to alarm pheromone. All statistics using Pearson’s Chi-Square. Asterisks or letters denote *posthoc* statistical significance (p < 0.05). Sample size per group, N = 18-29.

Surprisingly, we also observed that while young Day 1 bees are accepted by related guards, they are often rejected by unrelated guards (Figure 6B). This finding contradicts the broadly accepted “blank slate” hypothesis, which predicts that because day-old bees are devoid of any defining chemical signals, they should be always accepted by guards independent of relatedness (Breed et al., 2004). While we do not yet know which components of the CHC profile of young bees might have triggered a rejection by unrelated guards in our colonies, one plausible interpretation of these data is that the observed response of guards to unrelated Day 1 bees is an artifactual experimental outcome of a forced behavioral interaction between two bee groups, which in colonies with a typical demography, do not normally interact in the context of the hive entrance. These data suggested that nestmate recognition is a plastic behavior, that can be altered by social and other environmental contexts. To directly test this hypothesis, we next investigated the effects of exposing guard bees to the main component of alarm pheromone (isopentyl acetate), which is released by guards at the entrance to the colony when intruders are detected (Alaux and Robinson, 2007), and presented them with foraging nestmates. We found that in the context of a “high alert” state, guards reject nestmates at similar rates to rejections of non-nestmates under normal circumstances (Figure 6C, Pearson’s Chi-Squared, χ^2^ = 12.54, df = 2, p = 0.002). These data further indicate that nestmate recognition is a highly contextual behavior (Couvillon et al., 2013; Downs and Ratnieks, 2000).

## Discussion

The ability of colonies of social insects to reliably recognize group members is one of the remarkable adaptations that enabled their immense ecological success. Yet, how this complex task is accomplished remains unknown for many species. The data presented here indicate that different social insect species may have evolved different mechanistic solutions to their common problem of how to recognize group members. We show that in contrast to chemical gestalt mechanisms, which have been shown to play a role in some social insect species, nestmate recognition in colonies of the honey bee depends on a set of chemical cues that undergo regulated stereotypic developmental changes, and do not completely mature until bees make their final transition to foraging.

A major line of investigation in understanding nestmate recognition of social insects has been to determine how cue colony-specificity is determined. Cue specificity has historically been proposed to either be determined by mechanisms under genetic control or acquired from the environment (Crozier and Dix, 1979). Although our studies do not directly address the mechanism by which cue specificity is determined in honey bees, data from cross-fostering experiments suggest that cue development and specificity are defined by interactions between factors derived from the colony of origin of individual workers and the actual hive environment they live in. Therefore, our data suggest that, in contrast to being solely genetically determined or environmentally acquired, CHC profiles of honey bee workers develop via a biphasic process that is governed, at least in part, by the intrinsic physiology of individual workers, the specific behavioral tasks they are engaged in, and the hive environment they age in. In phase one, similar to other social insect species (Soroker et al., 1995), the total CHC amount builds up, possibly to increase the resistance of workers to desiccation while still inside the protective hive environment (Chung and Carroll, 2015). In phase two, the total amount of CHCs remains constant but the relative abundances of individual components shift in association with the behavioral maturation of workers, the worker’s colony environment, and probably environment- and age-dependent transcriptional changes in CHC biosynthetic enzymes.

Which specific components of the honey bee CHC profile specify nestmate recognition remains unknown. Although it has been shown that CHCs are likely the main cues used for nestmate recognition in honey bees (van Zweden and D’Ettorre, 2010), it is unlikely that all components of the CHC profile contribute to nestmate recognition (Akino et al., 2004; Dani et al., 2005, 2001; Martin et al., 2008; Ruther et al., 2002). In fact, it has previously been shown that alkenes likely play a more prominent role in nestmate recognition in the honey bee than alkanes (Dani et al., 2005). This is further supported by our data, which indicate that although unrelated foragers raised in the same colony are equally accepted, their overall CHC profiles remained somewhat qualitatively different (Figure 3C). These data provide two important insights. First, guards are not likely to use the full CHC profile of individuals to determine group membership. Second, differences in the CHC profiles of co-fostered foragers that originated from different source colonies further indicate that the cue maturation process does not follow the passive CHC transfer and homogenization that is characteristic of species that use the gestalt mechanism.

The lack of empirical support for a passive, “Gestalt”-like mechanism for the regulation of nestmate recognition cues in honey bees is further supported by our finding that the expression levels of genes that are known to play a role in CHC biosynthesis change in association with the observed developmental plasticity of the CHC profiles of workers. Although these data do not directly contradict the possibility of some passive transfer of particular CHCs, they do indicate that at least some components of the developing CHC profile are due to intrinsic plasticity in the transcriptional activity of the CHC synthesis pathway. However, our studies also importantly show that genetically-related bees that age in different colonies exhibit qualitatively different CHC profiles and CHC biosynthesis gene expression levels in their cuticles, which suggests that the intrinsic CHC maturation process could be also modulated by factors associated with the hive and/ or social environments. Based on these data, we propose a new model, which stipulates that the hive environment plays an indirect role in defining nestmate recognition cues by modulating the intrinsic pheromone synthesis pathway. Thus, the colony environment drives the development of similar pheromone profiles across all individuals living in the same colony, which in typical honey bee hives, seems to be directly associated with age-dependent division of labor. Such a model could resolve previous seemingly contradictory data which suggested that honey bee CHC profiles are defined by genetic (Page et al., 1991) versus environmental (Downs and Ratnieks, 1999) factors.

Previous studies suggested that the lower amounts of total CHCs in young bees represent a “blank slate” in terms of the nestmate recognition cue because these bees are readily “accepted” when introduced into unrelated colonies (Breed et al., 2004). In fact, this phenomenon was exploited here to introduce cohorts of bees to foster colonies, typically by placing the new bees on the top frames of experimental hives.Therefore, we were surprised by our observation that Day 1 bees are accepted at the entrance to their natal colony but rejected at the entrance of an unrelated colony. This apparent conundrum highlights an important, but often misunderstood, aspect of the nestmate recognition system in honey bees and other social insect species, which is that the “rejection” behavior by guards is highly contextual. Conceptually analogous to other biological systems that are responsible for detection of “self” versus “non-self” (*e.g.*, the acquired immunity system in vertebrates), behaviors associated with nestmate recognition are restricted to interactions between guards and incoming bees at the entrance to the hive (Couvillon et al., 2013). Therefore, because nestmate recognition is limited to a specific interaction between entering bees and guards at the entrance, the “acceptance” of day old bees outside the context of the hive’s entrance is actually due to the lack of a “rejection” behavior context within the hive.

Since Day 1 bees were rejected by an unrelated colony, yet accepted at their natal colony, it is possible that nestmate recognition of young bees either depends on components of the CHC profile that are already present in Day 1 bees, non-CHC chemicals, or an altogether different sensory modality. Alternatively, because newly eclosed bees usually perform cell cleaning behaviors at the interior of the hive, and therefore do not typically interact with guards at the hive entrance (Robinson, 1992; Smith et al., 2008; Søvik et al., 2015), differences in rejection of Day 1 bees between these two colonies might represent an experimental artifact resulting from differences in tolerance to the forced behavioral interaction between two bee groups that normally do not interact. Additionally, it has previously been shown that observed levels of guarding behaviors in honey bees are plastic, and could fluctuate in response to various environmental factors such as overall colony size, food availability, and “robbing” pressures from other colonies or predators (Downs and Ratnieks, 2000). Likewise, more extreme forms of plasticity in nestmate recognition systems have been reported in other social species. For example, some social insects can switch between using visual or chemosensory modalities for nestmate recognition under different circumstances (Baracchi et al., 2015). Furthermore, we found that when acutely exposed to the main component of alarm pheromone, guards will reject their nestmates, whom they would usually accept under normal circumstances (Figure 6C). These data clearly demonstrate that although chemosensory communication plays an important role in the nestmate recognition system of honey bees, cue specificity is not required for inducing rejection behaviors by guards, and that when presented with an acute “emergency” situation, acute alarm pheromone-dependent reduction in rejection threshold is more adaptive than selective rejection based on hive membership. Together, it seems that instead of being driven by simple binary decisions, nestmate recognition systems depend on plastic recognition of “friends” and “foes” as part of a broader group-level optimization of colony fitness that depends on social and other environmental contexts.

In conclusion, we propose that nestmate recognition cue production and acquisition in honey bees are not likely to be driven by processes that are analogous to the passive “Gestalt”-like mechanisms described in some ant species (Boulay et al., 2000; Lenoir et al., 2001; Meskali et al., 1995; Soroker et al., 1994; Victoria Soroker et al., 1995; Van Zweden et al., 2010). Instead, we propose that in honey bees, nestmate recognition cues, which are recognized and used by guards to define colony membership in the context of the entrance to the hive, mature in individual workers as integral elements of the overall CHC profile development via an intrinsic ontogenetic process that is associated with the stereotypic age-dependent behavioral maturation of workers, and is modulated by yet unknown social- and/or hive-specific factors. Altogether, data presented here suggest that during Hymenopteran evolution, different social species independently evolved analogous “solutions” for their common problem of how to recognize nestmates.

## Methods

### Animal Husbandry and Bee Collections

Honey bee (*Apis mellifera*) colonies were reared and managed using standard beekeeping techniques across two locations near St. Louis, MO: Tyson Research center (38° 31’N, 90° 33’W) and a residential home. For all experiments that included collections of bees at specific ages, capped brood frames were taken from a colony and placed in a humidified 32°C incubator. Once eclosed, about 1000 bees (<24 hours old) were marked with a spot of paint (Testors, Vernon Hills, IL, USA) on their thorax, and then reintroduced into either their natal or a foster colony, depending upon the experiment. For collections of bees at specific ages, marked bees were collected from internal frames of the colony one day post reintroduction (Day 1), seven days post reintroduction (Day 7), 14 days (Day 14) post reintroduction, and as returning foragers, identified by pollen loads on their hind legs or having a distended abdomen due to nectar loads, between 18 and 21 days post reintroduction. Bees used for chemical and molecular analyses were placed in individual 1.7 mL microtubes and immediately placed on dry ice. All samples were kept at -80°C until further analysis.

### Single-cohort colonies

Single-cohort colonies (SCC) were established as previously reported (Ben-Shahar et al., 2004, 2002; Greenberg et al., 2012; Whitfield et al., 2003). In short, about 1000 newly eclosed bees (<24 hours old) were placed in a small wooden nucleus hive-box with a young, unrelated mated queen, one honey frame from their natal colony, an empty comb frame, and three new frames with wax covered plastic foundation. Bees were collected as typical-aged nurses and precocious foragers one week after introduction, and as over-aged nurses and typical-aged foragers at three weeks after introduction. Bee samples were collected and stored as above.

### Undertaker collection

To induce “undertaking” behavior, about 1000 dead bees were placed into the top of two different colonies, and the first 20 bees that were observed removing dead bees from the colony were collected from the entrance. Returning foragers and in-hive nurses of unknown ages were also collected from each colony at the same time. Samples were stored and processed as described above.

### Cross-fostering experiment

1000 day-old bees from two independent source colonies were collected and marked as above. Half of the bees in each marked cohort were randomly reintroduced to both their own natal colony and the reciprocal foster colony. Subsequently, marked bees of defined age were recollected from internal frames of each colony as described above.

### Nestmate recognition assay

Every day over a three-week period, newly eclosed bees (<24 hours old) from a single source colony were collected as described above, uniquely color-marked, and then reintroduced into their natal colony. Subsequently, on each experimental day, bees from the following groups were collected, placed in individual 15 mL plastic tubes (Corning, Corning, NY, USA), and chilled on wet ice in an ice cooler up to 10 minutes before the assay in order to limit heat related stress: bees of the focal age (identified by color of mark), returning nectar foragers (denoted by distended abdomen and lack of pollen) of unknown age from the natal colony, and returning nectar foragers of unknown age from an unrelated colony. All foragers, which served as behavioral controls, were painted the same color as the experimental bees just after collection. Tubes were numbered in a randomized order and blinded to the experimenter conducting the behavioral assays. Fifteen bees per group were prepared for each colony each experimental day.

Behavioral assays were conducted simultaneously at two colonies (natal and unrelated) by two researchers, as well as recorded using digital video cameras. As described previously (D’Ettorre et al., 2006; Downs and Ratnieks, 2000), acceptance at the colony entrance was used as a proxy for nestmate recognition by placing individual bees on a modified entrance platform and recording the behavioral reactions of guard bees for ∼5 min. Bees were considered ‘Rejected’ if they were bit, stung and/or dragged by at least one guard bee (Video 1). Bees were considered ‘Accepted’ if they were approached by guards, antennated and/or licked and then left alone (not bit), if they immediately entered the colony and were not removed by other bees, or if they remained on the platform and did not receive aggression (Video 2). After 5 min, focal bees that remained on the platform outside the colony were removed before the next assay. All behaviors were scored in real time, and videos were retained as back-up. All behavioral assays were conducted during a period of 10 days, between 12 and 4pm, with two days focusing on each age of experimental bee (N = 20-30 bees per group).

Studies of the effect of alarm pheromone on nestmate recognition were done as above with the following modifications. First, guards were presented with nestmates or non-nestmates, collected as returning nectar foragers of unknown age, to assess baseline discriminatory behavior without pheromone, then were presented with nestmates in the presence of a filter paper containing 20 μl of 1:10 isopentyl acetate in mineral oil (Sigma Aldrich, St. Louis, MO USA) solution placed on the platform (Alaux and Robinson, 2007). To maintain constant pheromone exposure, 20 μl of solution was reapplied to the filter paper every 20 minutes over the course of the assays.

### Cuticular Hydrocarbon Extractions and GC analysis

CHCs were extracted from whole bees by placing individual bees into 6 mL glass vials fitted with 16mm PTFE/silica septa screw caps (Agilent Crosslab, Santa Clara, CA, USA). Bee CHCs were extracted in 500 uL hexane containing 10 ng/μl of octadecane (C_18_) and 10 ng/μl of hexacosane (C_26_), which served as injection standards. To achieve efficient extraction, each vial was gently agitated by vortexing (Fisher Scientific, Waltham, MA, USA) for 2 min at minimum speed. Extracts were immediately transferred to new 2 mL glass vials fitted with 9mm PTFE lined caps (Agilent Crosslab, Santa Clara, CA, USA). In cases where experiments involved forager honey bees, all bees (including non-foragers) had their hind legs removed prior to extraction, in order to ensure removal of pollen. 100 ul of each extract was transferred to a new 2 mL glass vial and stored at -20°C for further analysis; the remaining 400 uL was stored at -80°C as back-up.

Representative pooled samples of foragers and nurses of known age were first analyzed by combined gas chromatography/mass spectrometry (GC/MS) for compound identification. Samples were run from 150°(3 min hold) to 300°at 5°/min. Compounds were identified by their fragmentation pattern as compared to synthetic compounds. For profile characterizations of individual bees, samples were analyzed using an Agilent 7890A gas chromatograph system with a flame ionization detector (GC/FID) and PTV injector (cool-on-column mode), and outfitted with a DB-1 20 m x 0.18 mm Agilent 121- 1022 fused silica capillary column (Agilent Technologies, Inc. Santa Clara, CA, USA). Sample volumes of 1.0 μl were injected onto the column. Helium was the carrier gas and applied at a constant flow rate of 1 ml/min. Analysis of the extract was carried out with a column temperature profile that began at 50C (held for 1 min) and was ramped at 36.6 °C/min to 150C and then at 5C/min to 280C, where it was held for 10 min. The injector and FID temperatures were programmed to 280C and 300C, respectively. Agilent OpenLAB CDS (EZChrom Edition) software was used to calculate the retention time and total area of each peak.

### CHC Biosynthesis Gene Identification, RNA Isolation and Quantitative Real-Time PCR

Members of the highly conserved desaturase and elongase gene families were identified in the honey bee genome by using the protein BLAST search tool (https://blast.ncbi.nlm.nih.gov/Blast.cgi) with annotated *Drosophila melanogaster* amino acid sequences (https://flybase.org) of elongase and desaturase genes known to play a role in CHC biosynthesis (Chung and Carroll, 2015). Initial homologs in the honey bee genome were chosen by picking the top match (highest total score and query cover, lowest E value) for each *D. melanogaster* gene, and possible paralogs of these putative genes were identified by subsequently using the protein BLAST tool with these genes’ amino acid sequences. Many of these putative elongase and desaturase genes have previously been identified as possible CHC biosynthesis pathway genes in the honey bee (Falcón et al., 2014). To create phylogenetic trees, an LG model test (Le and Gascuel, 2008) was first performed with an estimative of invariable sites and a gamma heterogeneity rate between sites, for elongase and desaturase amino acid sequences separately, using MEGA 7 (Kumar et al., 2016). The –LogL (log likelihood) values were -4862.18 and -7202.65 for elongases and desaturases, respectively. Molecular phylogeny of both gene classes was performed using maximum likelihood analysis (Felsenstein, 1981) with 500 bootstrap replications. The *D. melanogaster* genes *EloF* and *desat1* were used as outgroups for the elongase and desaturase trees, respectively.

To measure mRNA levels of individual genes, the cuticles from the abdomens of four bees per group were dissected out, and total RNA was extracted using the Trizol Reagent (Life Technologies, Grand Island, NY, USA). SuperScript II (Life Technologies, Grand Island, NY, USA) reverse transcriptase was used to generate cDNA templates from 500ng of total RNA per sample by using random hexamers. A Bio-Rad (Hercules, CA, USA) CFX Connect Real-Time PCR Detection System and Bio-Rad iTaq Universal SYBR Green Supermix were subsequently used for estimating relative differences in mRNA levels across samples (N=4 per group, run in triplicate technical replications). The *eIF3-S8* housekeeping gene was used as a loading control as previously described (Alaux et al., 2009; Fischer and Grozinger, 2008; Greenberg et al., 2012; Mao et al., 2015). The specific primers for each gene are listed in Table 2.

### Statistical Analysis

All CHC analyses included a set of 19 peaks that represent well-established honey bee CHCs, identified by comparing GC traces to published data (Kather et al., 2011). For the comparisons of total CHCs across groups (as in Figure 1A), total ng of all identified CHCs in each bee were analyzed using ANOVA followed by Tukey’s HSD in R 3.3.2 (R Core Team, 2016). For the remainder of the datasets, the relative proportion of each compound in each sample was calculated and then used in further statistical analysis. For each dataset, a permutation MANOVA was run using the ADONIS function in the vegan package of R with Bray-Curtis dissimilarity measures (Oksanen et al., 2017). Pairwise comparisons with FDR p-value correction were subsequently run on experiments where more than two groups were compared. Data were visualized using non-metric multidimensional scaling (metaMDS function in the vegan package of R (Oksanen et al., 2017)) using Bray-Curtis dissimilarity, and either 2 or 3 dimensions in order to minimize stress to < 0.1. For Table 1 and Table 1 – supplement 1, an ANOVA followed by Tukey’s HSD post-hoc comparison, or Kruskal-Wallis followed by Dunn’s Test with FDR adjustment was performed using proportions of each compound across bees of the four time point collections. For behavioral data, the proportion of bees accepted by guard honey bees was calculated for each experimental group at each colony at each day of age. A Pearson’s chi-square was run for each day of age at each colony with subsequent pairwise comparisons. For qPCR data, relative expression levels were calculated as previously described (Greenberg et al., 2012; Hill et al., 2017; Zheng et al., 2014), using *eIF3-S8* as a loading control. Fold-expression data were generated by using the 2^-ΔΔCT^ method (Livak and Schmittgen, 2001) and designating a single individual from the “Day 1” group (Figure 5) as a calibrator. For statistical analyses, the 2^-ΔΔCT^ scores were compared within each gene across bees of different groups using an ANOVA followed by Tukey’s HSD post-hoc comparison, or Kruskal-Wallis followed by Dunn’s Test with FDR adjustment.

## Acknowledgements

We thank Tyson Research Center faculty and staff and Ellen Hartz for beekeeping assistance. We thank Anthony Cantu and Brice Henson for assistance in bee collections, Iris Chin for assistance with dissections and RNA extractions, and Kyle Skottke for hive building assistance.

---

**Figure 1 – figure supplement 1:**
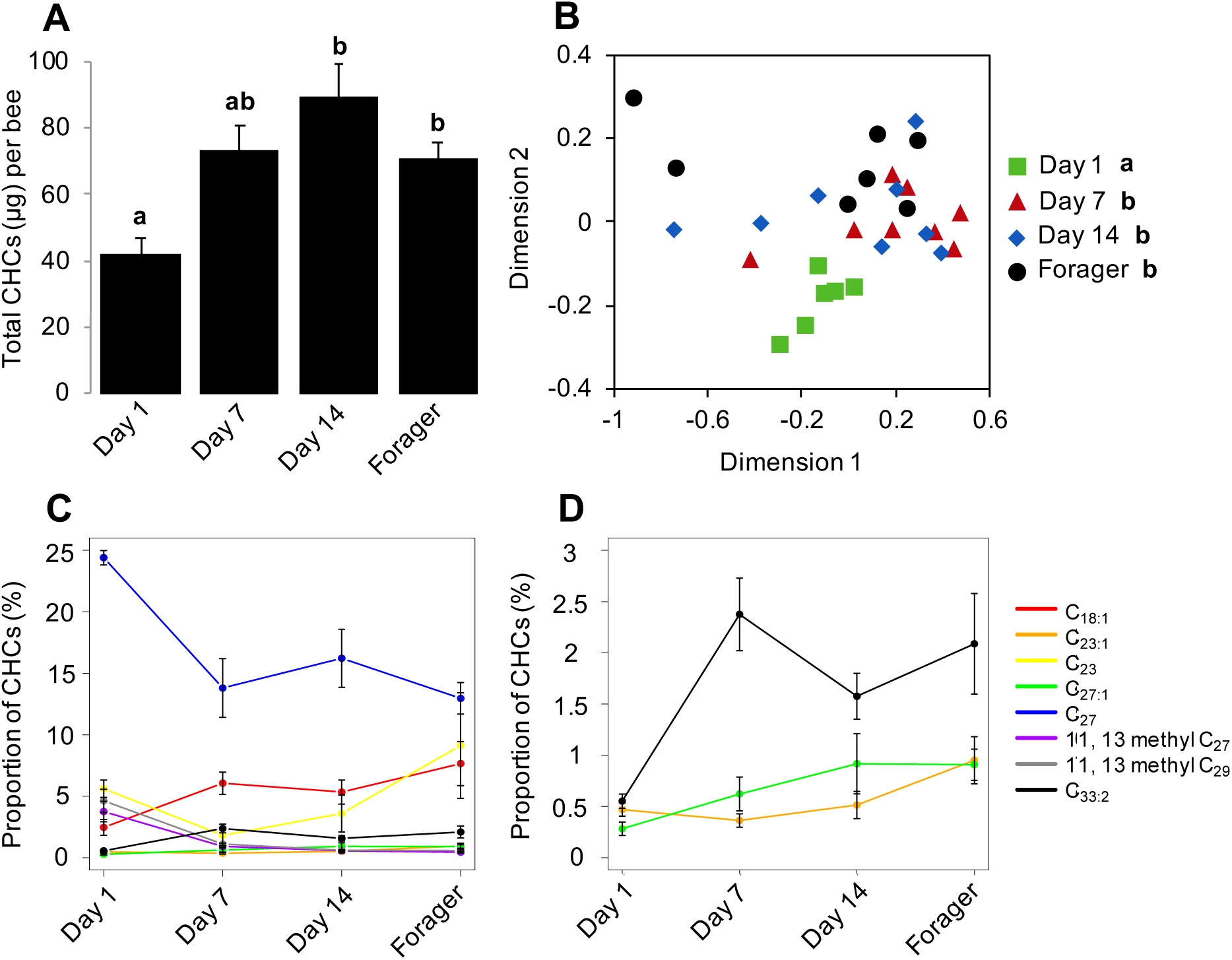
CHC profiles of bees exhibit quantitative and qualitative changes in association with age. Data shown here and in Fig. 1 were collected from two independent colonies. (A) Total CHC amounts (μg) extracted from sister bees of different ages. (B) CHC profiles of sister bees of different ages. (C) Statistically significantly changing proportions of individual CHCs across sister bees of different ages. D. A subset of C with low proportions. Statistics in A using ANOVA followed by Tukey’s HSD post-hoc. Statistics in B using Permutation MANOVA followed by FDR pairwise contrasts, shown as a non-metric multidimensional scaling plot depicting Bray-Curtis dissimilarity between samples. Statistics for C & D are listed in Table 1 – table supplement 1. Letters in graph and legend denote *posthoc* statistical significance (p < 0.05). Sample size per group, N = 8.

**Figure 5 – figure supplement 1:**
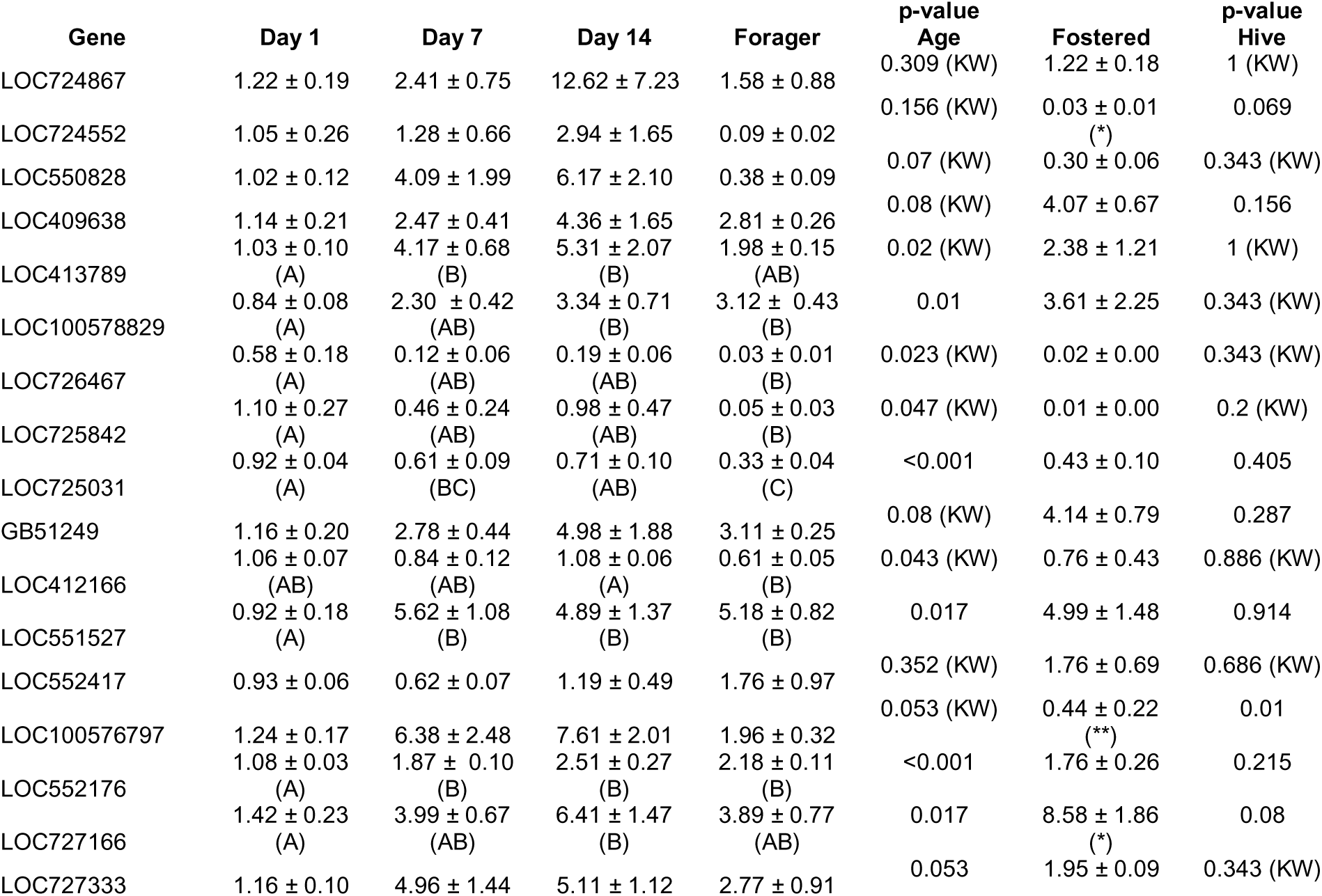
Genes differ in relative mRNA expression level between bees of different ages, and foraging sister bees raised in two different colonies. Numbers represent mean relative mRNA expression level ± standard error across four biological replicates. All p-values are from parametric ANOVA or nonparametric Kruskal Wallis ANOVA (denoted by “KW”). Letters denote statistically significant age groups across individual compounds via Tukey’s HSD (ANOVA post- hoc) or Dunn’s Test with FDR adjustment (KW post-hoc) (p < 0.05).

**Figure 5 – figure supplement 2:**
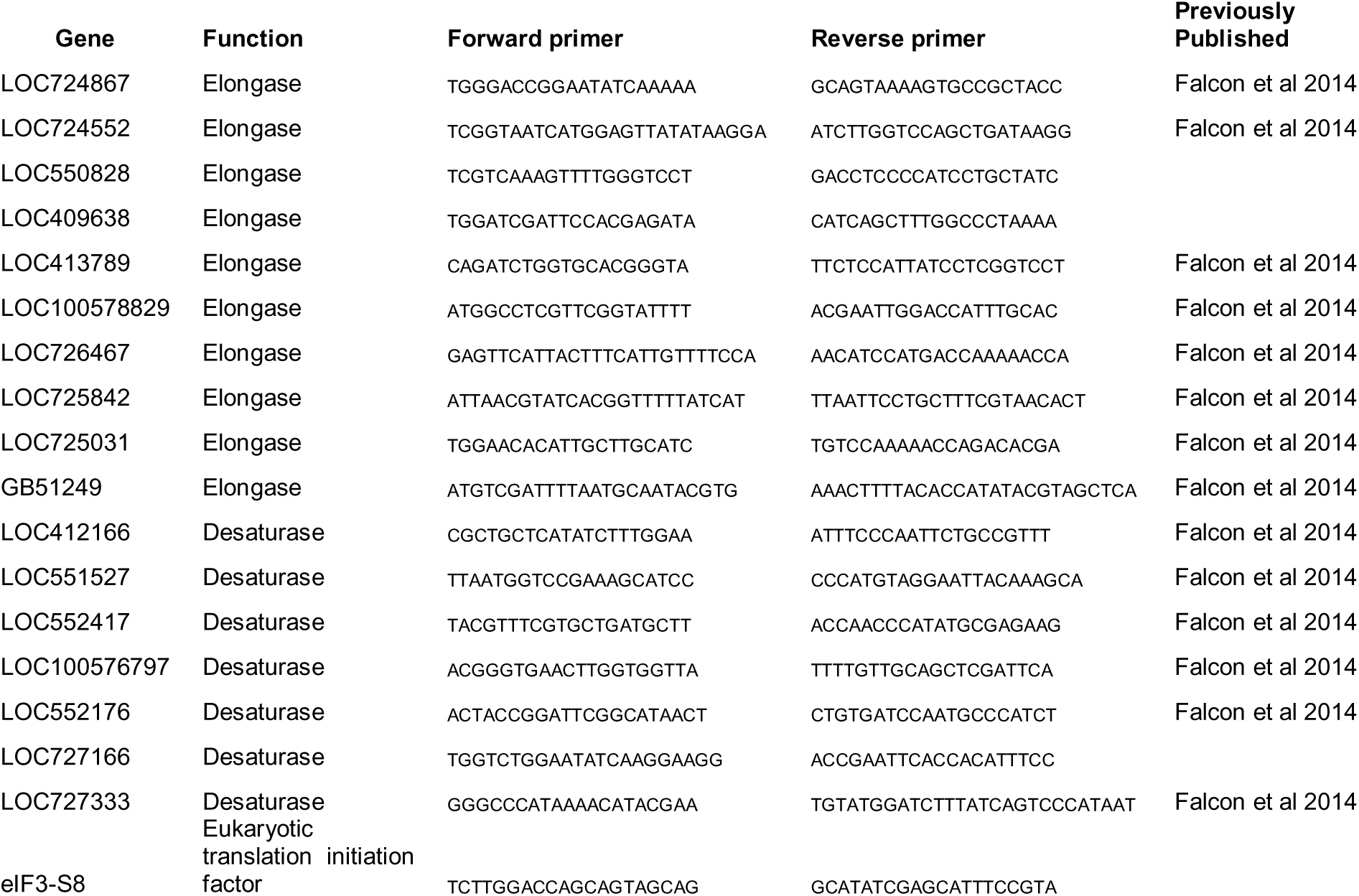
CHC biosynthesis genes and quantitative real- time PCR primers used in this study.

**Table S1.**
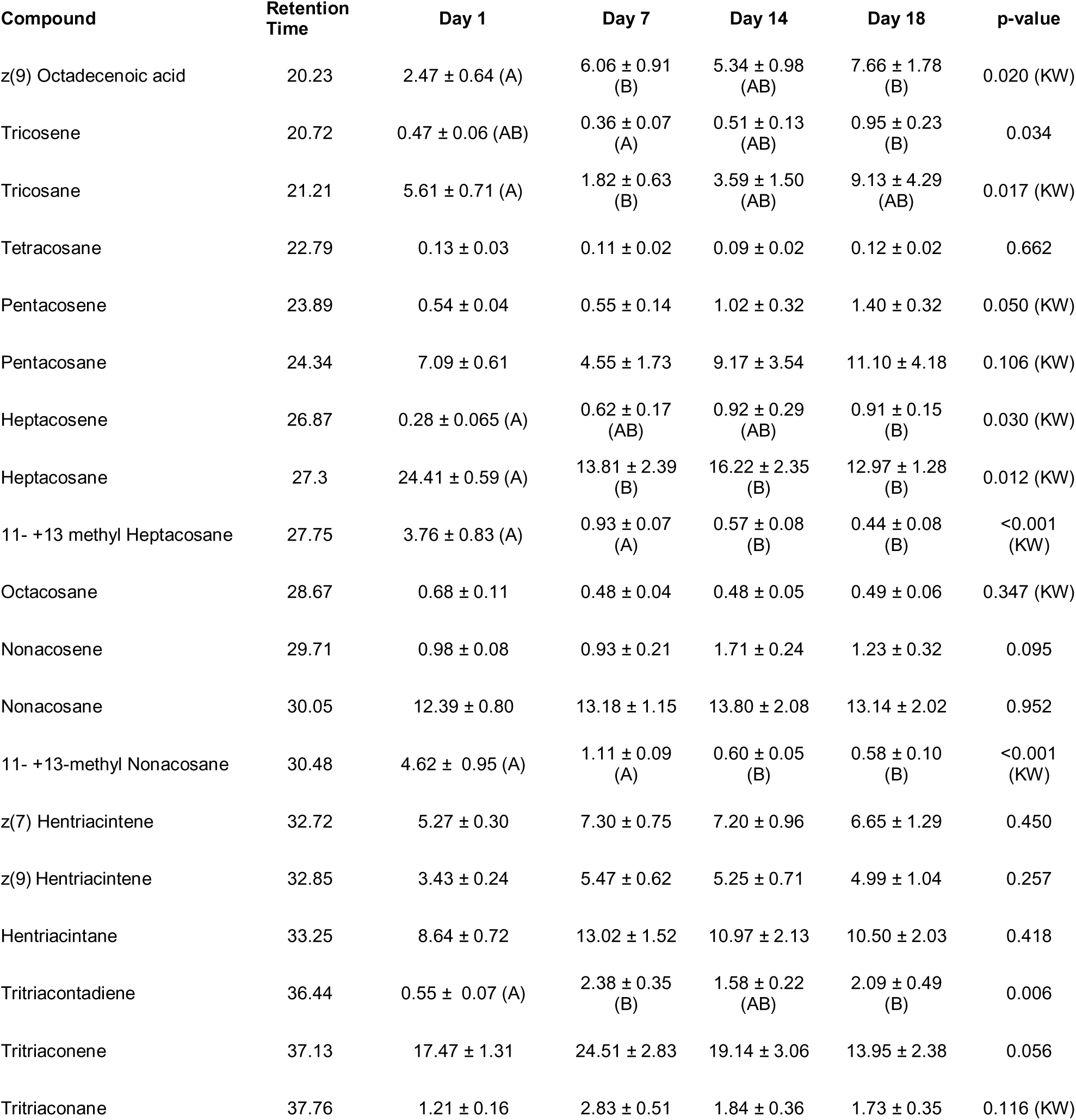
– table supplement 1: Individual compounds vary in proportion across different aged sister bees of a second colony. Numbers represent mean proportion of compound across bees of that age ± standard error. All p-values are from parametric ANOVA or nonparametric Kruskal Wallis ANOVA (denoted by “KW”). Letters denote statistically significant age groups across individual compounds via Tukey’s HSD (ANOVA post-hoc) or Dunn’s Test with FDR adjustment (KW post-hoc) (p < 0.05).

## Video Legends

**Video 1:** An interaction between a guard and focal bee scored as “Rejected”. The focal bee is marked with a green dot on its thorax.

**Video 2:** An interaction between a guard and focal bee scored as “Accepted”. The focal bee is marked with a pink dot on its thorax.

## References

Akino T, Yamamura K, Wakamura S, Yamaoka R. 2004. Direct behavioral evidence for hydrocarbons as nestmate recognition cues in Formica japonica (Hymenoptera: Formicidae). Appl Entomol Zool 39:381–387. doi:10.1303/aez.2004.38.

Alaux C, Robinson GE. 2007. Alarm Pheromone Induces Immediate–Early Gene Expression and Slow Behavioral Response in Honey Bees. J Chem Ecol 33:1346–1350. doi:10.1007/s10886-007-9301-.

Alaux C, Sinha S, Hasadsri L, Hunt GJ, Guzmán-Novoa E, DeGrandi-Hoffman G, Uribe-Rubio JL, Southey BR, Rodriguez-Zas S, Robinson GE. 2009. Honey bee aggression supports a link between gene regulation and behavioral evolution. Proc Natl Acad Sci U S A 106:15400–15405. doi:10.1073/pnas.090704310.

Baracchi D, Petrocelli I, Chittka L, Ricciardi G, Turillazzi S. 2015. Speed and accuracy in nest-mate recognition: A hover wasp prioritizes face recognition over colony odour cues to minimize intrusion by outsiders. Proc R Soc B Biol Sci 282:20142750. doi:10.1098/rspb.2014.275.

Ben-Shahar Y, Dudek NL, Robinson GE. 2004. Phenotypic deconstruction reveals involvement of manganese transporter malvolio in honey bee division of labor. J Exp Biol 207:3281–3288. doi:10.1242/jeb.0115.

Ben-Shahar Y, Robichon A, Sokolowski MB, Robinson GE. 2002. Influence of Gene Action Across Different Time Scales on Behavior. Science (80-) 296:741–744. doi:10.1126/science.106991.

Boehm T. 2006. Quality Control in Self/Nonself Discrimination. Cell. doi:10.1016/j.cell.2006.05.01.

Boulay R, Hefetz A, Soroker V, Lenoir A. 2000. Camponotus fellah colony integration: Worker individuality necessitates frequent hydrocarbon exchanges. Anim Behav 59:1127–1133. doi:10.1006/anbe.2000.140.

Breed MD. 1998. Recognition Pheromones of the Honey Bee. Bioscience 48:463–470. doi:10.2307/131324.

Breed MD, Cook CN, McCreery HF, Rodriguez M. 2015. Social Recognition in Invertebrates: The Honeybee as a Model System In: Aquiloni L, Tricarico E, editors. Social Recognition in Invertebrates. Cham, Switzerland: Springer International Publishing. pp. 147–164. doi:10.1007/978-3-319-17599-.

Breed MD, Perry S, Bjostad LB. 2004. Testing the blank slate hypothesis: Why honey bee colonies accept young bees. Insectes Soc 51:12–16. doi:10.1007/s00040-003- 0698-.

Breed MD, Williams KR, Fewell JH. 1988. Comb wax mediates the acquisition of nest- mate recognition cues in honey bees. Proc Natl Acad Sci U S A 85:8766–9. doi:10.1073/pnas.85.22.876.

Buckle GR, Greenberg L. 1981. Nestmate recognition in sweat bees (Lasioglossum zephyrum): Does an individual recognize its own odour or only odours of its nestmates? Anim Behav 29:802–809. doi:10.1016/S0003-3472(81)80014-.

Buczkowski G, Kumar R, Suib SL, Silverman J. 2005. Diet-related modification of cuticular hydrocarbon profiles of the argentine ant, Linepithema humile, diminishes intercolony aggression. J Chem Ecol 31:829–843. doi:10.1007/s10886-005-3547-.

Buczkowski G, Silverman J. 2006. Geographical variation in Argentine ant aggression behaviour mediated by environmentally derived nestmate recognition cues. Anim Behav 71:327–335. doi:10.1016/j.anbehav.2005.04.01.

Carlin NF, Hölldobler B. 1988. Influence of virgin queens on kin recognition in the carpenter ant Camponotus floridanus (Hymenoptera: Formicidae). Insectes Soc 35:191–197. doi:10.1007/BF0222393.

Carlin NF, Hölldobler B. 1987. The Kin Recognition System of Carpenter Ants (Camponotus spp.): II. Larger Colonies. Behav Ecol Sociobiol 20:209–217.

Carlin NF, Hölldobler B. 1986. The Kin Recognition System of Carpenter Ants (Camponotus spp.): I. Hierarchical Cues in Small Colonies Author (s): Norman F. Carlin and Bert Hölldobler Published by?: Springer Stable URL?: http://www.jstor.org/stable/4599936 REFERENCES Linked refere 19:123–134.

Carlin NF, Hölldobler B. 1983. Nestmate and Kin Recognition in Interspecific Mixed Colonies of Ants. Science (80-) 222:1027–1029.

Chung H, Carroll SB. 2015. Wax, sex and the origin of species: Dual roles of insect cuticular hydrocarbons in adaptation and mating. BioEssays 37:822–830. doi:10.1002/bies.20150001.

Couvillon MJ, Caple JP, Endsor SL, Kärcher M, Russell TE, Storey DE, Ratnieks FLW. 2007. Nest-mate recognition template of guard honeybees (Apis mellifera) is modified by wax comb transfer. Biol Lett 3:228–230. doi:10.1098/rsbl.2006.061.

Couvillon MJ, Segers FHID, Cooper-Bowman R, Truslove G, Nascimento DL, Nascimento FS, Ratnieks FLW. 2013. Context affects nestmate recognition errors in honey bees and stingless bees. J Exp Biol 216:3055–3061. doi:10.1242/jeb.08532.

Crozier RH, Dix MW. 1979. Analysis of two genetic models for the innate components of colony odor in social Hymenoptera. Behav Ecol Sociobiol 4:217–224. doi:10.1007/BF0029764.

D’Ettorre P, Wenseleers T, Dawson J, Hutchinson S, Boswell T, Ratnieks FLW. 2006. Wax combs mediate nestmate recognition by guard honeybees. Anim Behav 71:773–779. doi:10.1016/j.anbehav.2005.05.01.

Dani FR, Jones GR, Corsi S, Beard R, Pradella D, Turillazzi S. 2005. Nestmate Recognition Cues in the Honey Bee: Differential Importance of Cuticular Alkanes and Alkenes. Chem Senses 30:477–489. doi:10.1093/chemse/bji04.

Dani FR, Jones GR, Destri S, Spencer SH, Turillazzi S. 2001. Deciphering the recognition signature within the cuticular chemical profile of paper wasps. Anim Behav 62:165–171. doi:10.1006/anbe.2001.171.

Downs SG, Ratnieks F. 2000. Adaptive shifts in honey bee (Apis mellifera L.) guarding behavior support predictions of the acceptance threshold model. Behav Ecol 11:326–333. doi:10.1093/beheco/11.3.32.

Downs SG, Ratnieks FLW. 1999. Recognition of conspecifics by honeybee guards uses nonheritable cues acquired in the adult stage. Anim Behav 58:643–648. doi:10.1006/anbe.1999.117.

Errard C. 1994. Long-term memory involved in nestmate recognition in ants. Anim Behav. doi:10.1006/anbe.1994.124.

Espelie KE, Wenzel JW, Chang G. 1990. Surface lipids of social wasp Polistes melricus say and its nest and nest pedicel and their relation to nestmate recognition. J Chem Ecol 16:2229–2241. doi:10.1007/BF0102693.

Falcón T, Ferreira-Caliman MJ, Franco Nunes FM, Tanaka ÉD, do Nascimento FS, Gentile Bitondi MM. 2014. Exoskeleton formation in Apis mellifera: Cuticular hydrocarbons profiles and expression of desaturase and elongase genes during pupal and adult development. Insect Biochem Mol Biol 50:68–81. doi:10.1016/j.ibmb.2014.04.00.

Felsenstein J. 1981. Evolutionary trees from {DNA} sequences: a maximum likelihood approach. J Mol Evol 17:368–376.

Fischer P, Grozinger CM. 2008. Pheromonal regulation of starvation resistance in honey bee workers (Apis mellifera). Naturwissenschaften 95:723–729. doi:10.1007/s00114-008-0378-.

Gamboa GJ, Reeve HK, Ferguson ID, Wacker TL. 1986. Nestmate recognition in social wasps: the origin and acquisition of recognition odours. Anim Behav 34:685–695. doi:10.1016/S0003-3472(86)80053-.

Getz WM. 1982. An analysis of learned kin recognition in hymenoptera. J Theor Biol 99:585–597. doi:10.1016/0022-5193(82)90212-.

Getz WM. 1981. Genetically based kin recognition systems. J Theor Biol 92:209–226. doi:10.1016/0022-5193(81)90288-.

Greenberg JK, Xia J, Zhou X, Thatcher SR, Gu X, Ament SA, Newman TC, Green PJ, Zhang W, Robinson GE, Ben-Shahar Y. 2012. Behavioral plasticity in honey bees is associated with differences in brain microRNA transcriptome. Genes Brain Behav 11:660–70. doi:10.1111/j.1601-183X.2012.00782..

Hamilton WD. 1964a. The genetical evolution of social behaviour. I. J Theor Biol 7:1–16. doi:10.1016/0022-5193(64)90038-.

Hamilton WD. 1964b. The genetical evolution of social behaviour. II. J Theor Biol 7:17–52. doi:10.1016/0022-5193(64)90039-.

Hefetz A. 2007. The evolution of hydrocarbon pheromone parsimony in ants (Hymenoptera: Formicidae)—Interplay of colony odor uniformity and odor idiosyncrasy. A review. Myrmecological News 10:59–68.

Heinze J, Foitzik S, Hippert A, Hölldobler B. 1996. Apparent dear-enemy phenomenon and environment-based recognition cues in the ant Leptothorax nylanderi. Ethology 102:510–522. doi:10.1111/j.1439-0310.1996.tb01143..

Hill A, Zheng X, Li X, McKinney R, Dickman D, Ben-Shahar Y. 2017. The Drosophila Postsynaptic DEG/ENaC Channel ppk29 Contributes to Excitatory Neurotransmission. J Neurosci 37:3171–3180. doi:10.1523/JNEUROSCI.3850- 16.201.

Hölldobler B, Michener CD. 1980. Mechanisms of identification and discrimination in social Hymenoptera In: Markl H, editor. Evolution of Social Behavior: Hypotheses and Empirical Tests. Weinheim, West Germany: Verlag Chemie. pp. 35–58.

Huang ZY, Robinson GE. 1992. Honeybee colony integration: worker-worker interactions mediate hormonally regulated plasticity in division of labor. Proc Natl Acad Sci U S A 89:11726–11729. doi:10.1073/pnas.89.24.1172.

Kather R, Drijfhout FP, Martin SJ. 2011. Task Group Differences in Cuticular Lipids in the Honey Bee Apis mellifera. J Chem Ecol 37:205–212. doi:10.1007/s10886-011- 9909-.

Kumar S, Stecher G, Tamura K. 2016. MEGA7: Molecular Evolutionary Genetics Analysis Version 7.0 for Bigger Datasets. Mol Biol Evol 33:1870–1874. doi:10.1093/molbev/msw05.

Lacy RC, Sherman PW. 1983. Kin Recognition by Phenotype Matching. Am Nat 121:489–512.

Lahav S, Soroker V, Meer RKV, Hefetz A. 2001. Segregation of colony odor in the desert ant Cataglyphis niger. J Chem Ecol 27:927–943. doi:10.1023/A:101038291922.

Le SQ, Gascuel O. 2008. An improved general amino acid replacement matrix. Mol Biol Evol 25:1307–1320. doi:10.1093/molbev/msn06.

Lenoir A, Cuisset D, Hefetz A. 2001. Effects of social isolation on hydrocarbon pattern and nestmate recognition in the ant Aphaenogaster senilis (Hymenoptera, Formicidae). Insectes Soc 48:101–109. doi:10.1007/PL0000175.

Liang D, Silverman J. 2000. “You are what you eat": diet modifies cuticular hydrocarbons and nestmate recognition in the Argentine ant, Linepithema humile. Naturwissenschaften 87:412–416. doi:10.1007/s00114005075.

Livak KJ, Schmittgen TD. 2001. Analysis of Relative Gene Expression Data Using Real- Time Quantitative PCR and the 2 ? ?? C T Method. Methods 25:402–408. doi:10.1006/meth.2001.126.

Makki R, Cinnamon E, Gould AP. 2014. The Development and Functions of Oenocytes. Annu Rev Entomol 59:405–425. doi:10.1146/annurev-ento-011613-162056

Mao W, Schuler MA, Berenbaum MR. 2015. A dietary phytochemical alters caste- associated gene expression in honey bees. Sci Adv 1:e1500795:1–9. doi:10.1126/sciadv.150079.

Martin SJ, Correia-Oliveira ME, Shemilt S, Drijfhout FP. 2018. Is the Salivary Gland Associated with Honey Bee Recognition Compounds in Worker Honey Bees (Apis mellifera)? J Chem Ecol. doi:10.1007/s10886-018-0975-.

Martin SJ, Helanterä H, Drijfhout FP. 2008. Colony-specific hydrocarbons identify nest mates in two species of Formica ant. J Chem Ecol 34:1072–1080. doi:10.1007/s10886-008-9482-.

Meskali M, Provost E, Bonavita-Cougourdan A, Clément JL. 1995. Behavioural effects of an experimental change in the chemical signature of the ant Camponotus vagus (Scop.). Insectes Soc 42:347–358. doi:10.1007/BF0124216.

Oksanen J, Blanchet FG, Friendly M, Kindt R, Legendre P, McGlinn D, Minchin PR, O’Hara RB, Simpson GL, Solymos P, Henry M, Stevens H, Szoecs E, Wagner H. 2017. vegan: Community Ecology Package.

Page RE, Metcalf RA, Metcalf RL, Erickson EH, Lampman RL. 1991. Extractable hydrocarbons and kin recognition in honeybee (Apis mellifera L.). J Chem Ecol 17:745–56. doi:10.1007/BF0099419.

Pusey A, Wolf M. 1996. Inbreeding avoidance in animals. Trends Ecol Evol 11:201–206. doi:10.1016/0169-5347(96)10028-.

Reeve HK. 1989. The Evolution of Conspecific Acceptance Thresholds. Am Nat 133:407–435.

Richard F-J, Errard C, Hefetz A, Christides J-P. 2004. Food influence on colonial recognition and chemical signature between nestmates in the fungus-growing ant Acromyrmex subterraneus subterraneus. Chemoecology 14:9–16. doi:10.1007/s00049-003-0251-.

Richard FJ, Poulsen M, Hefetz A, Errard C, Nash DR, Boomsma JJ. 2007. The origin of the chemical profiles of fungal symbionts and their significance for nestmate recognition in Acromyrmex leaf-cutting ants. Behav Ecol Sociobiol 61:1637–1649. doi:10.1007/s00265-007-0395-.

Robinson GE. 1992. Regulation of Division of Labor in Insect Societies. Annu Rev Entomol 37:637–665. doi:10.1146/annurev.en.37.010192.00322.

Ruther J, Sieben S, Schricker B. 2002. Nestmate recognition in social wasps: Manipulation of hydrocarbon profiles induces aggression in the European hornet. Naturwissenschaften 89:111–114. doi:10.1007/s00114-001-0292-.

Singer TL, Espelie KE. 1996. Nest surface hydrocarbons facilitate nestmate recognition for the social wasp, Polistes metricus say (Hymenoptera: Vespidae). J Insect Behav 9:857–870. doi:10.1007/BF0220897.

Smith CR, Toth AL, Suarez A V., Robinson GE. 2008. Genetic and genomic analyses of the division of labour in insect societies. Nat Rev Genet 9:735–748. doi:10.1038/nrg242.

Soroker V, Hefetz A, Cojocaru M, Franke S, Francke W. 1995. Structural and chemical ontogeny of the postpharyngeal gland in the desert ant Cataglyphis niger. Physiol Entomol 20:323–329.

Soroker V, Vienne C, Hefetz A. 1995. Hydrocarbones dynamics within and between nestmates in Cataglyphis niger (Hymenoptera: formicidae). J Chem Ecol 21:365–378. doi:doi:10.1007/BF0203672.

Soroker V, Vienne C, Hefetz A, Nowbahari E. 1994. The postpharyngeal gland as a “Gestalt” organ for nestmate recognition in the ant Cataglyphis niger. Naturwissenschaften 81:510–513. doi:10.1007/BF0113268.

Søvik E, Bloch G, Ben-Shahar Y. 2015. Function and evolution of microRNAs in eusocial Hymenoptera. Front Genet 6:1–11. doi:10.3389/fgene.2015.0019.

Stuart RJ. 1988. Collective cues as a basis for nestmate recognition in polygynous leptothoracine ants. Proc Natl Acad Sci U S A 85:4572–4575. doi:full text at www.pnas.or.

Team RC. 2016. R: A language and environment for statistical computing.

Teseo S, Lecoutey E, Kronauer DJC, Hefetz A, Lenoir A, Jaisson P, Châline N. 2014. Genetic Distance and Age Affect the Cuticular Chemical Profiles of the Clonal Ant Cerapachys biroi. J Chem Ecol 40:429–438. doi:10.1007/s10886-014-0428-.

Trivers RL. 1971. The Evolution of Reciprocal Altruism. Q Rev Biol 46:35–57.

Tsutsui ND. 2004. Scents of self?: The expression component of self / non- self recognition systems. Ann Zool Fennici 41:713–727.

Van Zweden JS, Brask JB, Christensen JH, Boomsma JJ, Linksvayer TA, d’Ettorre P. 2010. Blending of heritable recognition cues among ant nestmates creates distinct colony gestalt odours but prevents within-colony nepotism. J Evol Biol 23:1498–1508. doi:10.1111/j.1420-9101.2010.02020..

van Zweden JS, D’Ettorre P. 2010. Nestmate recognition in social insects and the role of hydrocarbons In: Blomquist GJ, Bagneres A-G, editors. Insect Hydrocarbons. Cambridge: Cambridge University Press. pp. 222–243. doi:10.1017/CBO978051171190.

Wagner D, Tissot M, Gordon D. 2001. Task-Related Environment Alters the Cuticular Hydrocarbon Composition of Harvester Ants. J Chem Ecol 27:1805–1819.

West SA, Griffin AS, Gardner A. 2007. Evolutionary explanations for cooperation. Curr Biol CB 17:R661–672. doi:10.1016/j.cub.2007.06.00.

Whitfield CW, Cziko A-M, Robinson GE. 2003. Gene Expression Profiles in the Brain Predict Behavior in Individual. Science (80-) 302:296–299. doi:10.1126/science.108680.

Wilkinson GR. 1988. Reciprocal altruism in bats and other mammals. Ethol Sociobiol 9:85–100.

Yew JY, Chung H. 2015. Insect pheromones: An overview of function, form, and discovery. Prog Lipid Res 59:88–105. doi:10.1016/j.plipres.2015.06.00.

Zheng X, Valakh V, Diantonio A, Ben-shahar Y, Louis S, States U. 2014. Natural antisense transcripts regulate the neuronal stress response and excitability 1–16. doi:10.7554/eLife.01849.

